# Host Metabolic Reprogramming Defines Macrophage Antifungal Activity Against *Cryptococcus neoformans*

**DOI:** 10.64898/2026.09.30.755789

**Authors:** Ayesha S. Nair, Dikshanta V. Luitel, Karen L. Wozniak

## Abstract

Innate immune cells, including macrophages and DCs, are critical for controlling infection by *Cryptococcus neoformans*. Although *C. neoformans* is phagocytosed equally by each pulmonary phagocyte subset, CD11b^+^ DCs support fungal growth, whereas Ly6c^-^ monocyte-like macrophages are antifungal. The metabolic programs underlying these contrasting responses remain incompletely understood, but our previous work points to increased glycolysis in antifungal cells. Therefore, we hypothesized that metabolic reprogramming in phagocytes following cryptococcal exposure leads to different outcomes. We examined ten metabolic pathways using flow cytometry, metabolic inhibition, antifungal assays, and pulmonary infection. Cryptococcal exposure increased GLUT-1, HK, HIF-1α, and lactate production in J774A.1 macrophages, consistent with enhanced glycolytic metabolism in antifungal cells. In contrast, ATP5a and IDH2 were reduced, suggesting diminished mitochondrial metabolism. *Cryptococcus* also decreased G6PD, ACAC, ASS1, and PRDX2, indicating alterations in PPP activity, fatty acid synthesis, arginine metabolism, and antioxidant defense. Pharmacologic inhibition of PPP further reduced G6PD expression and significantly decreased intracellular cryptococcal burden, supporting a role for glucose-derived metabolic pathways in regulating fungal survival. In vivo, infection increased GLUT-1 and HK expression in antifungal female Ly6c^-^ monocyte-like macrophages but not in permissive CD11b^+^ DCs, whereas other metabolic markers showed limited or subset-specific changes. Collectively, these findings demonstrate that antifungal cells use a predominantly glycolytic program accompanied by reduced mitochondrial, anabolic, and antioxidant-associated protein expression following interaction with *C. neoformans*. This phagocyte subset-dependent metabolic response during pulmonary infection may influence antifungal activity and identify metabolic pathways that could be targeted to strengthen host defense.

## INTRODUCTION

*Cryptococcus neoformans* is an encapsulated fungal pathogen responsible for pulmonary and disseminated infections, with severe disease frequently manifesting as cryptococcal meningitis. Individuals with HIV/AIDS are at greatest risk, although infection is also observed among organ transplant recipients, and individuals undergoing immunosuppressive treatment [1]. Cryptococcal infection continues to be a global health burden, accounting for about 112,000 annual deaths worldwide [2]. Infection is generally initiated following inhalation of dessicated yeast or cryptococcal spores. Within the lungs, *C. neoformans* encounters a heterogenous population of innate immune cells such as macrophages and dendritic cells. These innate immune cells recognize and internalize cryptococcal cells and can promote fungal clearance through several antimicrobial mechanisms [3]. However, *C. neoformans* can also survive and replicate within certain phagocyte populations, thereby facilitating dissemination from the lungs to other organs, including the central nervous system [4–6]. Therefore, understanding the cellular mechanisms that determine whether phagocytes restrict or support cryptococcal growth is important for identifying new strategies to improve antifungal immunity.

An emerging factor influencing the outcome of host-pathogen interactions is the metabolic response of immune cells [7,8]. Activation of immune cells is accompanied by changes in metabolic states, thereby providing energy to support immune functions [9]. Interaction of innate immune cells with pathogens leads to metabolic shifts in immune cells [10]. One such response that is observed in a lot of infections is the alterations in glucose metabolism. For example, previous studies in our lab have shown increased glucose dependence when antifungal immune cells interact with *C. neoformans* [11]. This has also been shown with immune cell responses to other fungal and bacterial pathogens such as *Aspergillus fumigatus, Mycobacterium tuberculosis (M. tb)* and *Candida albicans* [12–14]. These observations suggest that increased glucose utilization may represent a common metabolic adaptation that supports antimicrobial functions of antifungal phagocytes.

A shift toward increased glycolysis can resemble the Warburg effect [15]. In activated immune cells, this allows rapid utilization of glucose and provides intermediates required for immune effector functions [16,17]. However, glucose metabolism does not function independently of other metabolic pathways. Glucose-derived intermediates can also enter the pentose phosphate pathway (PPP), which generates NADPH required for redox regulation and NADPH dependent processes such as ROS production. Mitochondrial pathways, including the tri carboxylic acid cycle (TCA cycle) and oxidative phosphorylation (OXPHOS), as well as the fatty acid oxidation pathway (FAO) can also undergo substantial changes during immune cell activation [18–20]. Together, these interconnected pathways can determine the metabolic and functional state of innate immune cells during infection.

Our previous studies suggest that glucose metabolism is important in the immune response to *C. neoformans.* Using SCENITH, we previously demonstrated that increased dependence on glycolytic pathways in J774A.1 macrophages and Ly6c^-^ monocyte-like macrophages is essential for antifungal immunity [11]. However, mechanisms underlying this increased glucose dependence and contribution of glucose metabolism to macrophage antifungal activity remain incompletely understood. Furthermore, whether cryptococcal exposure alters additional metabolic pathways that could contribute to the functional state of macrophages has not been fully characterized.

In the present study, we investigated metabolic changes in innate immune cells following exposure to *C. neoformans.* Based on our previous observation of increased glucose dependence, we hypothesized that cryptococcal exposure would be accompanied by increased expression of proteins involved in glucose uptake and glycolytic metabolism. Additionally, we also evaluated proteins associated with PPP, mitochondrial metabolism, antioxidant responses, fatty acid synthesis, etc, to characterize broader metabolic changes occurring during cryptococcal exposure. Together, this study will expand our understanding of the metabolic adaptations of innate immune cells during cryptococcal infection and provides insight into the relationship between host cell metabolism and antifungal activity.

## MATERIALS AND METHODS

### Fungal strains

*Cryptococcus neoformans* strains H99 (serotype A, mating type α) and mCherry producing strain JLCN920 derived from the KN99 background (serotype A, mating type α), a kind gift of Dr. Jennifer Lodge (Duke University, Durham, NC) were used in this study. The strains were maintained as 15% glycerol stocks at -80°C. Before each experiment, the fungal strains were cultured in yeast peptone dextrose (YPD) broth (BD Difco; Franklin Lakes, NJ) for 18h at 30°C in a shaking incubator. The cells were subsequently collected, washed 3 times with sterile phosphate buffered saline (PBS) and enumerated using a hemocytometer. Trypan blue exclusion dye was used to assess cell viability.

### Cell culture

The murine macrophage cell line J774A.1 was obtained from ATCC (TIB-67; ATCC; Manassas, VA). Cells were maintained in Dulbecco’s modified eagle medium (DMEM; Thermo Fisher Scientific, Waltham, MA) supplemented with 10% heat-inactivated fetal bovine serum (FBS), 10% NCTC-109, 1% non-essential amino acids, 100U penicillin/ml, and 100 µg streptomycin/ml, then filter-sterilized through a 0.22µm filter. J774A.1 macrophages were cultured in CytoOne T75 tissue culture flasks (USA Scientific, Ocala, FL) at 37 °C, 5% CO_2_. Cells were passaged every 3-4 days according to manufacturer’s directions.

### Mice

Male and female BALB/c mice, 5-6 weeks of age, were obtained from Charles River Laboratories and housed in the Oklahoma State University Animal Resources facility. All experimental procedures were performed in accordance with guidelines set by Oklahoma State University’s Institutional Animal Care and Use Committee.

### Murine pulmonary cryptococcal infection

Male and female BALB/c mice (n=4 per group) were anesthetized with 2% isoflurane and intranasally inoculated with 1 × 10^4^ CFU of the KN99 mCherry strain suspended in 50 µL of sterile PBS. Control mice received 50ul of sterile PBS alone. At 3 days post-infection, mice were euthanized and lungs were aseptically harvested. The left lung lobes were homogenized in sterile PBS and used to determine fungal burden by CFU analysis. The right lung lobes were minced and enzymatically digested with collagenase IV (1 mg/mL; Sigma-Aldrich) in RPMI media (Thermo Fisher Scientific, Waltham, MA) at 37°C for 30 mins. The digested tissue was filtered through 70 and 40 µm nylon filters to obtain single-cell suspensions. Cells were washed with sterile Hank’s Balanced Salt Solution, and erythrocytes were removed with 1X red blood cell lysis buffer (eBioscience, San Diego, CA). The remaining cells were counted using a hemocytometer with trypan blue exclusion and were used for flow cytometric analysis.

### Flow cytometric analysis of surface & intracellular metabolic markers

J774A.1 macrophages were seeded at a concentration of 4×10^6^ cells/ml and incubated with KN99 mCherry strain at a ratio of 20:1 and 1ug/ml of anti-GXM antibody F12D2 (a kind gift of Dr. Tom Kozel, University of Nevada Reno, Reno, NV) for 30 mins, 1h or 24h to evaluate early and late metabolic responses respectively at 37°C and 5% CO_2_. Uninfected J774A.1 macrophages were included as controls. Following incubation, cells were centrifuged at 3000 rpm (2194 x g) for 3 mins at 4°C and resuspended in 200ul FACS buffer (2% fetal bovine serum in PBS, sterile filtered). 1µl Fc block (CD16/CD32) was added, followed by staining with surface antibodies such as CD45 and GLUT-1 along with staining for extracellular fungal organisms with calcofluor white (Thermo Fisher Scientific). For identifying pulmonary phagocyte populations, cells were stained with surface antibodies such as CD45, CD11c, F4/80, MHC-II, Ly6c, CD103 and CD11b as previously described [6,11]. To identify metabolic pathways involved in anti-cryptococcal activity from cell lines and primary cells, met-flow was conducted [21]. Cells were incubated with the surface flow antibodies along with GLUT-1 metabolic marker for 30 mins at 4°C. Post incubation, cells were centrifuged 3 times at 3000 rpm (2194g) for 3 mins at 4°C and then subjected to intracellular staining. Cells were permeabilized and stained intracellularly with anti-hexokinase (HK), anti-glucose-6-phosphate dehydrogenase (G-6PD), anti-ATP5a, anti-isocitrate dehydrogenase (IDH2), anti-hypoxia inducible factor (HIF-α), anti-peroxiredoxin (PRDX2), anti-acetyl CoA carboxylase, and anti-arginosuccinate synthase (ASS1) for 30 mins at room temperature. This was followed by staining with a secondary antibody conjugated with a fluorophore for 30 mins. Cells were then washed 3 times with permeabilization buffer. Samples were run on a NovoCyte 3000 flow cytometer (Agilent Technologies, Santa Clara, CA) and analyzed by NovoExpress software (Agilent).

### Lactate assay

J774A.1 macrophages were incubated with *C. neoformans* strain H99 at 20:1 ratio in the presence of 1µg/ml of anti-GXM antibody and incubated for 2h at 37°C and 5% CO_2_. Following infection, cells were treated with 2-deoxy-D-glucose (100mM), oligomycin (1µM), or a combination of both inhibitors for 25 mins. Macrophages treated with 1% DMSO were included as the control group. Following treatment, intracellular lactate levels were measured using a commercially available L-Lactate assay kit (Abcam, Waltham, MA) according to the manufacturer’s instructions. Colorimetric absorbance was measured at an optical density of 570nm using Synergy HTX multi-mode plate reader (BioTek).

### Inhibition of the pentose phosphate pathway

J774A.1 macrophages were seeded in 96-well plate at a concentration of 4×10^6^ cells/ml. The pentose phosphate pathway was inhibited using 6-aminonicotinamide (6-AN; Sigma Aldrich, St. Louis, MO), an inhibitor of G-6PD at a final concentration of 200µM/ well. DMSO treated macrophages were included as the control group. The cells were incubated for 24h at 37°C and 5% CO_2_. Following treatment, cells were washed with sterile PBS, and G-6PD protein expression was assessed by flow cytometry (as described above).

### ROS assay

To assess ROS production, untreated J774A.1 cells, antifungal J774A.1 cells exposed to H99, and treated with 6-AN, 2-DG, or DMSO were stained with CellROX reagent (ThermoFisher Scientific), a fluorogenic probe used to detect oxidative stress. Cells were incubated with the reagent at for 30 mins at 37°C and 5% CO_2_. Cells were then washed 3 times with PBS and analyzed by flow cytometry (excitation/emission: 485/520 nm).

### Quantitative real-time PCR (qRT-PCR)

J774A.1 cells were incubated alone or with the cryptococcal strain H99 and treated with either 6-AN, 2-DG, or DMSO for 24h. Following incubation, the cells were centrifuged at 3500 rpm (2986g) for 3 minutes at 4°C and resuspended in Trizol reagent overnight. Total RNA was isolated using the RNeasy mini kit (Qiagen, West Chester, PA), followed by cDNA synthesis using the Verso cDNA synthesis kit (ThermoFisher Scientific). The resulting cDNA was used to assess the expression of *Nox2* and *p47phox* by qRT-PCR. Taqman primers and TaqMan gene expression master mix (ThermoFisher Scientific) were used, with GAPDH as the housekeeping gene. Each reaction was prepared in a final volume of 20ul consisting of 19ul of the appropriate TaqMan reaction mixture and 1ul of cDNA. The samples were run on QuantStudio™ 5 Real-Time PCR System, 96-well, 0.2 mL machine (Applied Biosystems, Waltham, MA). Data was calculated using the 2^-ΔΔCt^ mathematical formula and analyzed on Microsoft Excel.

### Statistical Analysis

Statistical analyses were performed using GraphPad Prism version 5.0 (GraphPad Software). Comparison between two independent groups were performed using an unpaired, two-tailed *t* test with a 95% CI and *p < .05* Comparison involving three or more groups were performed using one-way analysis of variance followed by Tukey’s post test, with significant differences defined as *p < .05*.

## RESULTS

### J774A.1 macrophages undergo a glycolytic shift following cryptococcal infection, characterized by increased GLUT-1 and hexokinase expression

To define the metabolic requirements of antifungal and permissive cells during cryptococcal infection, we previously performed SCENITH analysis and observed increased glucose dependence in antifungal cells and reduced glucose dependence in permissive cells, indicating that host glycolysis is important for antifungal activity [11]. Thus, to further characterize the components underlying these metabolic differences, we evaluated key transporters and enzymes involved in glycolysis. Since glucose uptake is the first limiting step, we assessed expression of the glucose transporter GLUT-1. We observed a significant increase in the percentage of GLUT-1 expressing J774A.1 macrophages following treatment with KN99 mCherry cryptococcal strain compared with uninfected macrophages (Fig. 1A). We next evaluated the expression of hexokinase (HK), which catalyzes the first step of glycolysis by phosphorylating glucose to generate glucose-6-phosphate. Consistent with the GLUT-1 response, the percentage of HK expressing J774A.1 macrophages was significantly increased following treatment with KN99 mCherry cryptococcal strain compared with uninfected macrophages (Fig. 1B). Together, these findings support our previously published SCENITH results [11] and demonstrate that the increased glucose dependence of antifungal macrophages is accompanied by enhanced expression of proteins that facilitate glucose uptake and its entry into the glycolytic pathway.

**Fig 1:**
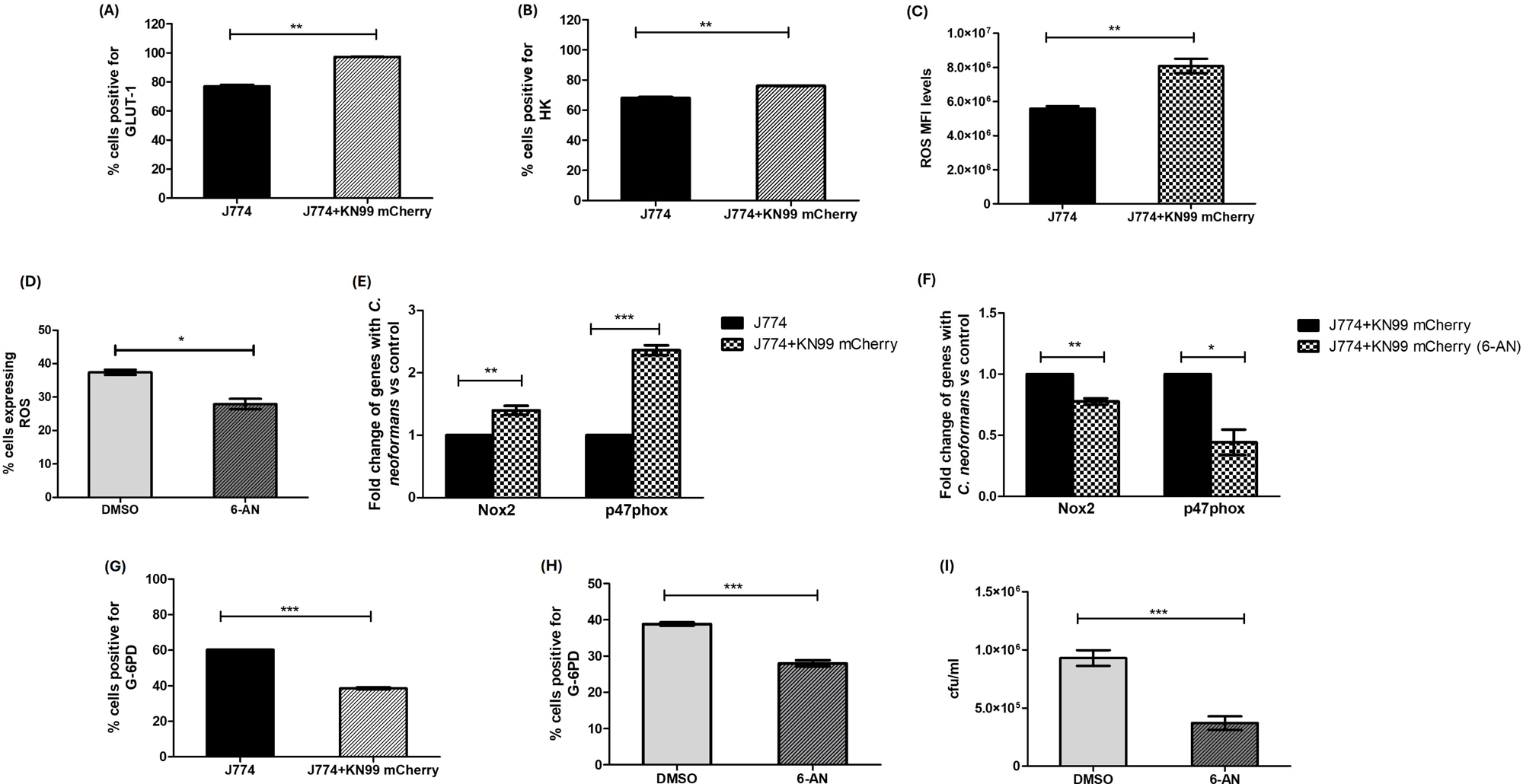
Cryptococcal exposure alters glucose metabolism and ROS production in J774A.1 macrophages, while 6-AN treatment reduces ROS, oxidase gene expression, and fungal burden. J774A.1 macrophages were incubated with KN99 mCherry strain at a ratio of 20:1 at 37°C and 5% CO_2_. Following incubation, cells were stained for surface and intracellular metabolic markers and analyzed by flow cytometry. Protein expression is presented as the percentage of cells positive for indicated metabolic marker. (A) Glucose transporter GLUT-1 expression was measured after 30 mins of cryptococcal uptake (B) Hexokinase (HK) expression was measured after 1h of cryptococcal uptake. (C) Intracellular ROS was measured by flow cytometry after 2h of cryptococcal uptake and is represented as mean fluorescent intensity (MFI). (D) J774A.1 macrophages were treated with 6-AN for 24h, after which intracellular ROS was measured by flow cytometry. (E) mRNA expression of *Nox2* and *p47phox* genes was measured after 24h of cryptococcal exposure and is shown as fold change relative to untreated J774A.1 macrophages. (F) *Nox2* and *p47phox* mRNA expression was measured in H99 treated macrophages with or without 200µm 6-AN. (G) Glucose-6-phosphate dehydrogenase (G6PD) expression was measured after 24h of cryptococcal uptake. (H) To assess the effect of PPP inhibition, J774A.1 macrophages were treated with 6-AN for 24h, and G6PD protein expression was subsequently evaluated by flow cytometry. DMSO treated macrophages were included as controls. (I) To determine the effect of PPP inhibition on antifungal activity, 6-AN treated macrophages were incubated with cryptococcal cells for 24h. Macrophages were lysed using sterile water, followed by plating the fungal cells on YPD agar for CFU enumeration. Data shown are means ± standard errors of the means (SEM) from three independent experiments (n=3). A two-tailed unpaired t-test with a 95% confidence interval (*p < 0.05*) was used to determine statistically significant differences.

### Cryptococcal exposure enhances ROS production and *Nox2/p47phox* expression in J774A.1 macrophages, both of which are reduced by PPP inhibition

Previous studies have linked increased glucose metabolism to ROS production in J774A.1 macrophages, leading us to examine ROS levels following *C. neoformans* exposure [22]. H99 exposed antifungal J774A.1 macrophages showed increased intracellular ROS compared with macrophages alone (Fig. 1C). To determine whether the increase in ROS depended on glycolysis, we measured intracellular ROS in J774A.1 macrophages after treatment with glycolytic inhibitor 2-deoxy glucose (2-DG). However, treatment with 2-DG did not significantly alter ROS levels (Fig. S4). Because increased glycolytic metabolism can provide glucose-6-phosphate for the pentose phosphate pathway (PPP), we next examined whether PPP contributed to ROS production in H99-exposed antifungal macrophages. Treatment with the PPP inhibitor 6-aminonicotinamide (6-AN) significantly reduced intracellular ROS levels in these cells (Fig. 1D). Since PPP generates NADPH used by NOX2 to produce ROS, we measured the mRNA expression of *Nox2* and *p47phox*, which encode the oxidase and a protein involved in its activation, respectively. H99 treatment increased the mRNA expression of *Nox2* and *p47phox* compared with macrophages alone (Fig. 1E), whereas 6-AN treatment reduced the expression of both genes in H99 treated cells (Fig. 1F). Together, these findings suggest that PPP contributes to the ROS response in H99 treated macrophages and that 6-AN treatment is associated with reduced *Nox2* and *p47phox* expression.

### J774A.1 macrophages restrict cryptococcal survival following inhibition of the pentose phosphate pathway

We next investigated the contribution of the PPP to the macrophage response during cryptococcal infection. We first assessed the expression of glucose-6-phosphate dehydrogenase (G6PD), the rate limiting enzyme of oxidative PPP. Interestingly, the percentage of G6PD positive J774A.1 macrophages was significantly decreased following treatment with KN99 mCherry cryptococcal strain compared with uninfected macrophages (Fig. 1G). To determine the functional significance of reduced PPP activity, J774A.1 macrophages were treated with 6-AN. Flow cytometric analysis showed that 6-AN further reduced the percentage of G6PD positive macrophages compared with the DMSO-vehicle control. (Fig. 1H). We subsequently assessed cryptococcal survival by quantifying fungal burden after macrophage treatment with 6-AN. Treatment with 6-AN significantly reduced cryptococcal CFUs compared with DMSO treated macrophages (Fig. 1I), indicating that PPP contributes to the regulation of intracellular cryptococcal survival and suggests that inhibition of this pathway promotes antifungal activity.

### J774A.1 macrophages exhibit reduced ATP5a and IDH2 expression following cryptococcal infection

In addition to increased glucose dependence, our previous SCENITH analysis revealed reduced mitochondrial dependence in antifungal cells, suggesting diminished reliance on oxidative phosphorylation (OXPHOS) [11]. To further characterize this metabolic shift, we examined ATP5a, a core subunit of mitochondrial ATP synthase that contributes to ATP production through OXPHOS. The percentage of ATP5 expressing J774A.1 macrophages was significantly reduced following infection with cryptococcal cells compared to macrophages alone (Fig. 2A). We next assessed mitochondrial isocitrate dehydrogenase 2 (IDH2), a NADP+ dependent enzyme that catalyzes the conversion of isocitrate to α-ketoglutarate in the tricarboxylic acid (TCA) cycle and contributes to mitochondrial redox homeostasis through NADPH production. Similar to ATP5a, the percentage of IDH2 expressing J774A.1 macrophages was significantly reduced following infection with cryptococcal cells compared to macrophages alone (Fig. 2B). Together, these findings support our previously published SCENITH results [11] and indicate that the increased glucose dependence of antifungal macrophages is accompanied by reduced expression of proteins associated with mitochondrial metabolism.

**Fig 2:**
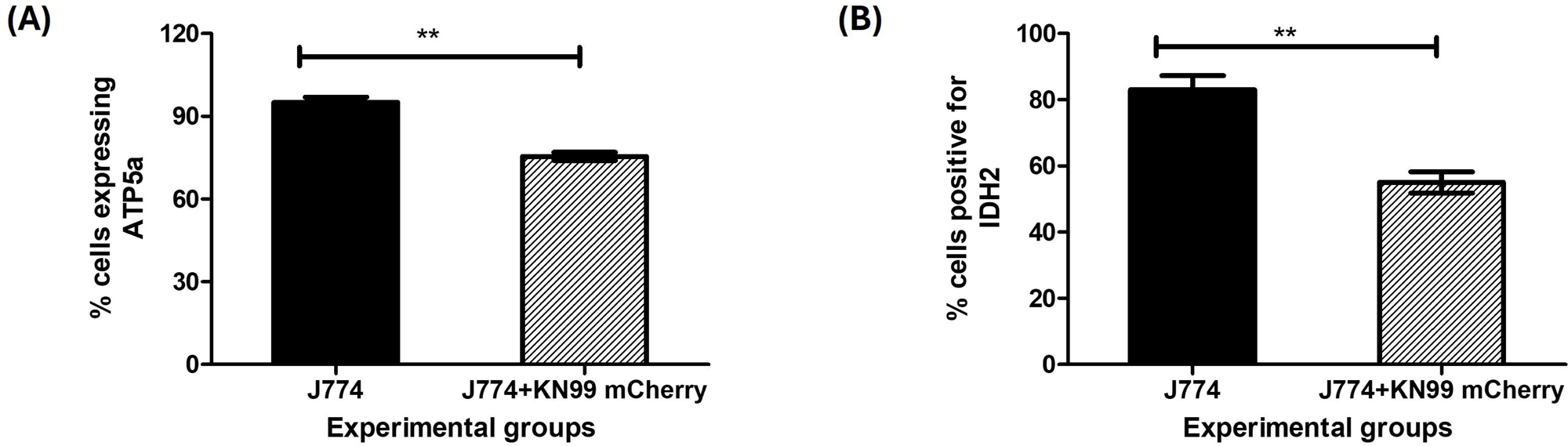
J774A.1 macrophages exhibit reduced expression of TCA cycle and OXPHOS associated proteins ATP5a and IDH2 expression respectively, following cryptococcal infection: J774A.1 macrophages were incubated with KN99 mCherry strain at a ratio of 20:1 at 37°C and 5% CO_2_. Following incubation, cells were stained with the surface marker anti-CD45 and intracellular antibodies against ATP5a and isocitrate dehydrogenase (IDH2), followed by flow cytometric analysis. Protein expression is presented as the percentage of cells positive for indicated metabolic marker. (A) ATP5a and (B) IDH2 protein expression was measured in J774A.1 macrophages following 24h of cryptococcal infection. Data shown are means ± standard errors of the means (SEM) from three independent experiments (n=3). A two-tailed unpaired t-test with a 95% confidence interval (*p < 0.05*) was used to determine statistically significant differences.

### Glycolytic reprogramming in antifungal cells during *Cryptococcus* interaction is lactate dependent and associated with HIF-1α activation (Warburg effect)

Given that antifungal immunity was associated with a metabolic shift from oxidative phosphorylation toward increased glucose utilization, we next investigated whether this resembles the Warburg effect, a metabolic signature often observed in cancer cells [15]. The Warburg effect is characterized by increased glucose uptake even in the presence of oxygen and enhanced lactate production, allowing rapid energy generation [23]. To assess this, we measured lactate accumulation in J774A.1 macrophages following exposure to *C. neoformans.* Infected macrophages exhibited significantly higher lactate levels compared to uninfected controls, consistent with a shift toward glycolysis. To determine whether this effect was dependent on glycolysis, cells were treated with 2-deoxy glucose to inhibit glycolysis. Under these conditions, the difference in lactate accumulation between infected and uninfected macrophages was not significant. In contrast, treatment with oligomycin, which inhibits only oxidative phosphorylation, restored the phenotype, resulting in elevated levels of lactate in H99 infected cells. Importantly, *C. neoformans* with and without metabolic inhibitor treatment showed no significant changes in lactate accumulation, indicating that the observed changes were due to host cell metabolic reprogramming rather than fungal metabolism (Fig. 3A). As the Warburg effect is known to be regulated by HIF-1α, we next assessed HIF-α protein expression by flow cytometry. We observed that the percentage of HIF-1α expressing J774A.1 macrophages was significantly increased following infection with cryptococcal cells compared to macrophages alone, further supporting the induction of Warburg effect during infection (Fig. 3B).

**Fig 3:**
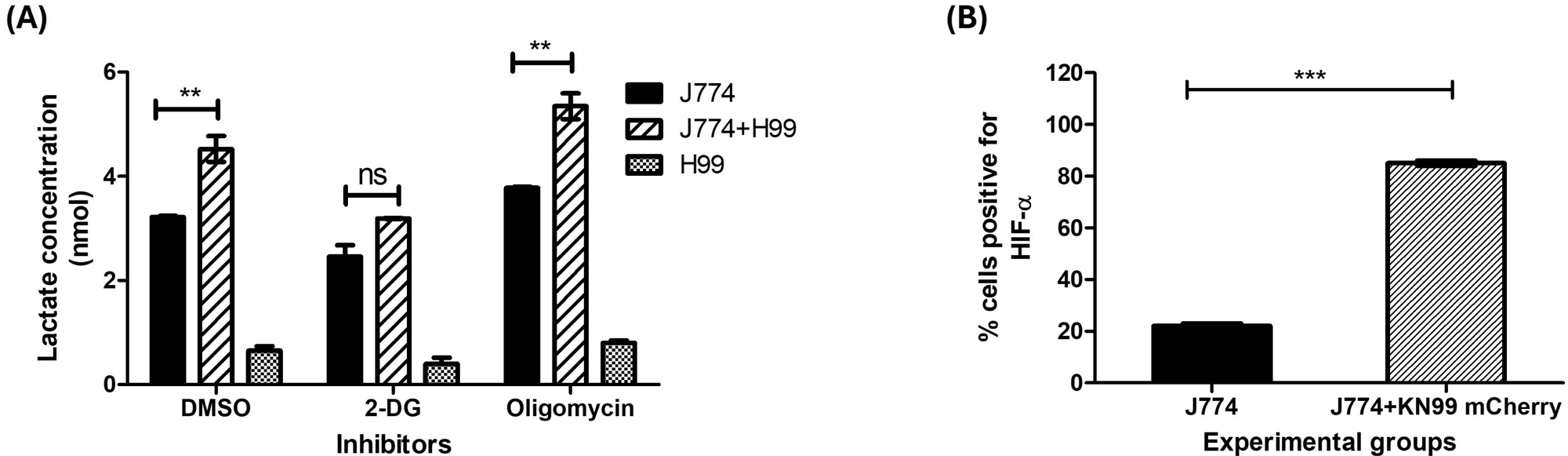
***C. neoformans* infection promotes Warburg- like metabolic reprogramming in J774A.1 macrophages:** (A) J774A.1 macrophages were incubated with KN99 mCherry strain at a ratio of 20:1 for 2h at 37°C and 5% CO_2_. Following uptake, cells were treated with the metabolic inhibitors such as 2-deoxy-D-glucose (2-DG), oligomycin, or a combination of both inhibitors for 25 mins. Cells were subsequently lysed, and intracellular lactate levels were measured using a colorimetric assay at an absorbance of 570nm. (B) After 1h of cryptococcal uptake by J774A.1 macrophages, intracellular HIF-α protein expression was measured by flow cytometry. Data shown are means ± standard errors of the means (SEM) from two independent experiments (n=2). One-way ANOVA followed by Tukey’s Multiple Comparison Test and a two-tailed unpaired t-test with a 95% confidence interval was performed, with significant differences indicated at *p < 0.05*.

### J774A.1 macrophages exhibit reduced expression of proteins associated with fatty acid and arginine biosynthesis following cryptococcal infection

We next investigated whether cryptococcal infection alters the anabolic capacity of J774A.1 macrophages by examining proteins involved in fatty acid synthesis and arginine metabolism. We first assessed acetyl CoA carboxylase (ACAC) which catalyzes the conversion of acetyl-CoA to malonyl-CoA, the rate limiting step in de novo fatty acid synthesis. Fatty acid synthesis supports lipid-mediated signaling, and the production of inflammatory lipid mediators required for immune cell function [24]. We observed that the percentage of ACAC expressing J774A.1 macrophages was significantly reduced following infection with cryptococcal cells compared to macrophages alone (Fig. 4A).

**Fig 4:**
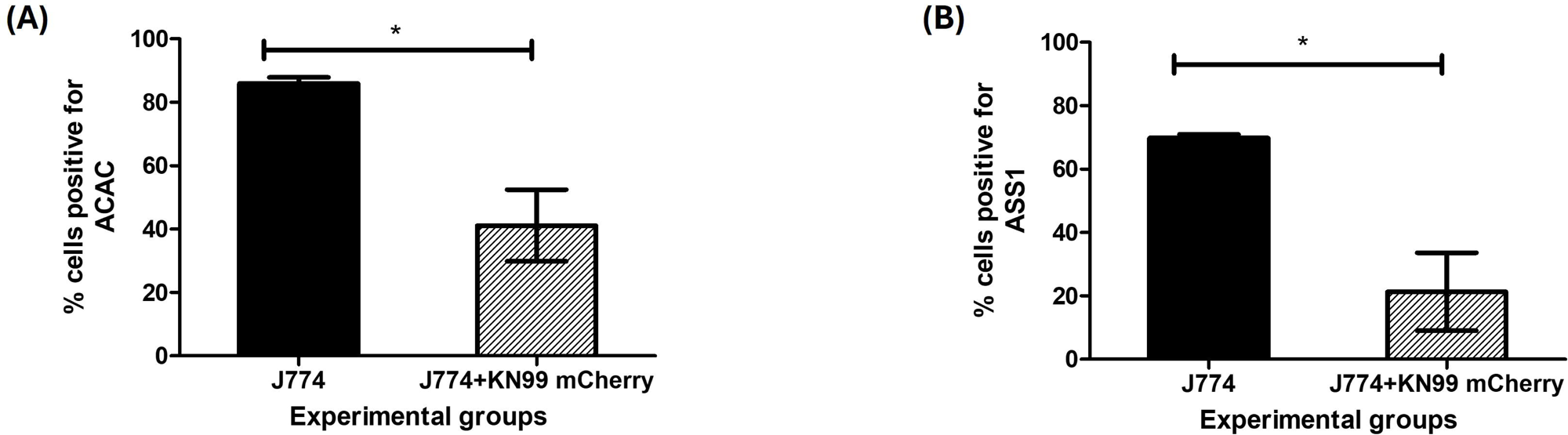
Anabolic reprogramming in J774A.1 macrophages during *C. neoformans* infection is characterized by reduced fatty acid synthesis and arginine metabolism associated proteins: J774A.1 macrophages were incubated with KN99 mCherry strain at a ratio of 20:1 at 37°C and 5% CO_2_. Following incubation, cells were stained with the surface marker anti-CD45 and intracellular antibodies against acetyl CoA carboxylase (ACAC) and argininosuccinate synthetase (ASS1), followed by flow cytometric analysis. Protein expression is presented as the percentage of cells positive for indicated metabolic marker. (A) Intracellular ACAC abundance and (B) ASS1 protein expression was measured in J774A.1 macrophages following 24h of cryptococcal infection. Data shown are means ± standard errors of the means (SEM) from three independent experiments (n=3). A two-tailed unpaired t-test with a 95% confidence interval (*p < 0.05*) was used to determine statistically significant differences.

We next examined argininosuccinate synthetase (ASS1) which catalyzes the formation of argininosuccinate from citrulline and aspartate and represents a key step in endogenous arginine biosynthesis. In macrophages, arginine availability influences several immune functions, including nitric oxide production. Similar to ACAC, the percentage of ASS1 expressing J774A.1 macrophages was significantly reduced following infection with cryptococcal cells compared to macrophages alone (Fig. 4B). Together, these findings indicate that cryptococcal infection is associated with reduced expression of proteins involved in fatty acid synthesis and arginine metabolism, consistent with a broader shift away from anabolic metabolism in J774A.1 macrophages.

### J774A.1 macrophages exhibit reduced antioxidant PRDX2 expression during cryptococcal infection

Because our metabolic analyses indicated altered mitochondrial activity during cryptococcal infection, we next examined peroxiredoxin 2 (PRDX2), an antioxidant enzyme that reduces hydrogen peroxide and contributes to cellular redox homeostasis. We observed that the percentage of PRDX2 expressing J774A.1 macrophages was significantly reduced following infection with cryptococcal cells compared to macrophages alone (Fig. 5). This finding is consistent with the increased intracellular ROS observed in H99 treated macrophages (Fig. 1C). Reduced PRDX2 expression may limit peroxide scavenging and contribute to a more oxidative intracellular environment that is potentially favorable for antifungal activity.

**Fig 5:**
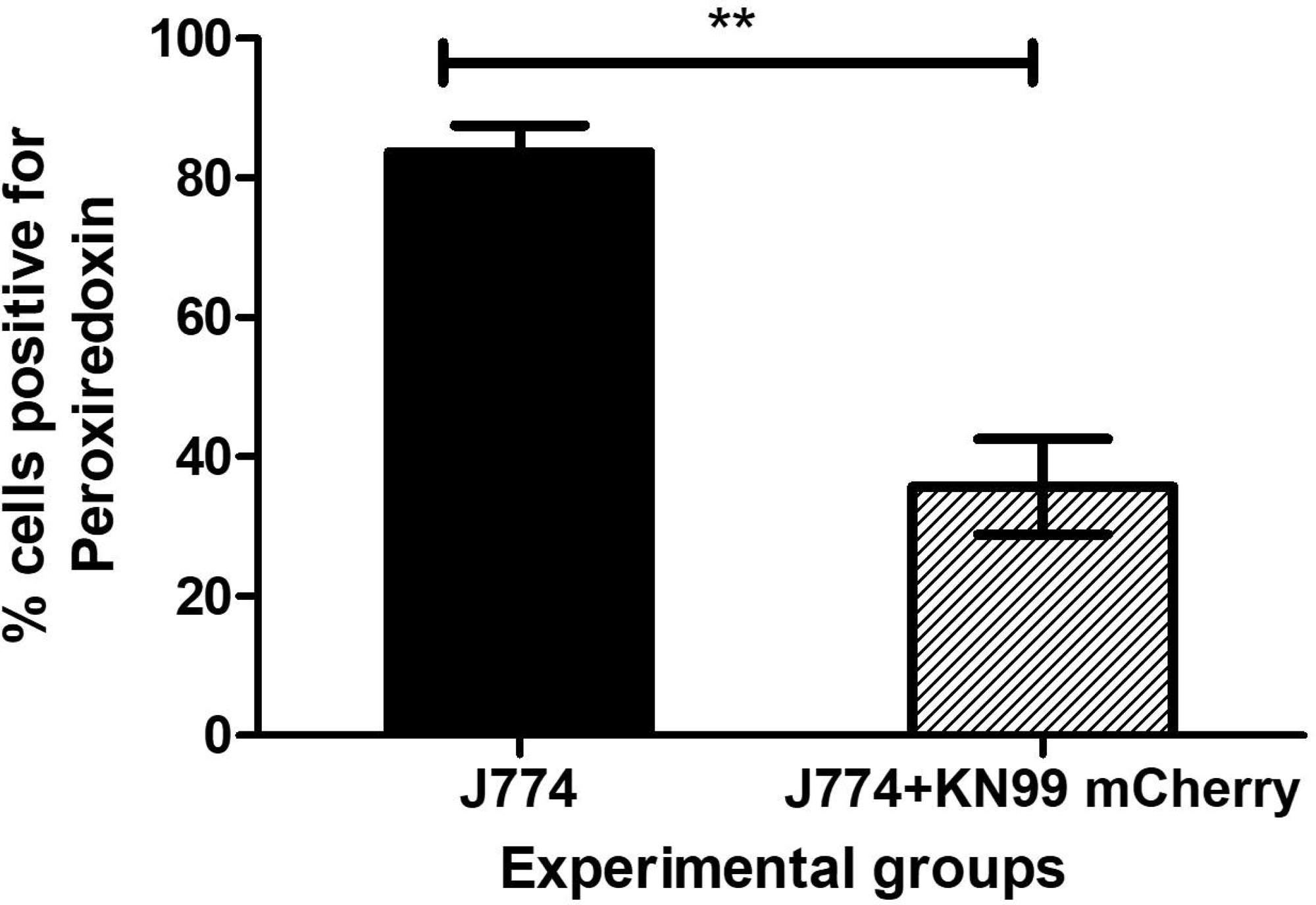
Altered antioxidant capacity in *C. neoformans* infected J774A.1 macrophages is characterized by PRDX2 downregulation: J774A.1 macrophages were incubated with KN99 mCherry strain at a ratio of 20:1 for 24h at 37°C and 5% CO_2_. Following incubation, cells were stained with the surface marker anti-CD45 and intracellular antibodies against peroxiredoxin (PRDX2), followed by flow cytometric analysis. PRDX2 expression is presented as the percentage of PRDX2-positive cells. Data shown are means ± standard errors of the means (SEM) from three independent experiments (n=3). A two-tailed unpaired t-test with a 95% confidence interval (*p < 0.05*) was used to determine statistically significant differences.

### Cryptococcal infection induces phagocyte subset-dependent metabolic changes in the lung

Following our *in vitro* studies in J774A.1 macrophages, we investigated whether cryptococcal infection induces similar metabolic changes in pulmonary phagocytes *in vivo.* Male and female Balb/c mice were infected intranasally with *C. neoformans* strain H99, and lungs were collected at 3 days post infection. Pulmonary leukocytes were analyzed by flow cytometry to identify alveolar macrophages (AMs), interstitial macrophages (IMs), CD11b^+^ DCs, CD103^+^ DCs, Ly6c^+^ and Ly6c^-^ monocyte-like macrophages using the gating strategy as described in Fig. S3. The expression of the metabolic and redox-associated proteins GLUT-1, HK, ATP5a, IDH2, G6PD, ACAC, ASS1 and PRDX2 were evaluated across these pulmonary phagocyte subsets.

Our previous SCENITH analysis demonstrated higher glucose dependence in female Ly6c^-^monocyte-like macrophages than in male CD11b^+^ DCs suggesting that glycolytic metabolism is preferentially associated with the antifungal subset [6,11]. Consistent with the response observed in J774A.1 macrophages, the percentage of GLUT-1 and HK positive female Ly6c^-^ monocyte-like macrophages were significantly increased following *C. neoformans* infection compared with uninfected controls (Fig. 6A). In contrast, infection did not significantly alter the percentages of GLUT-1 and HK positive male CD11b^+^ DCs (Fig. 6B). These findings reveal distinct glycolytic responses between the two pulmonary phagocyte subsets, with increases in GLUT-1 and HK characterizing antifungal Ly6c^-^ monocyte-like macrophages but not permissive CD11b^+^ DCs.

**Fig 6:**
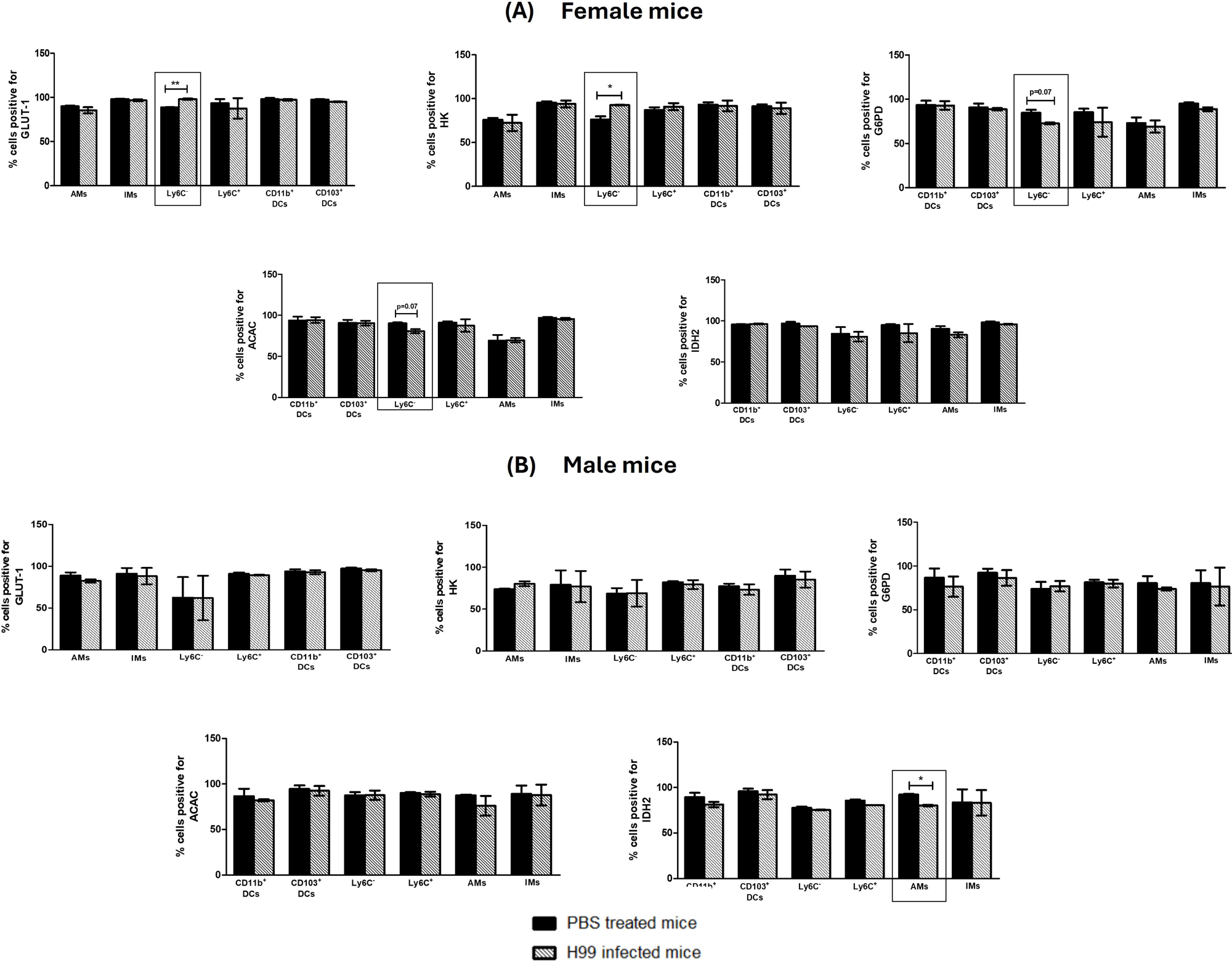
Expression of proteins associated with glycolysis, the pentose phosphate pathway, OXPHOS, and fatty acid synthesis in male and female pulmonary phagocyte subsets during *C. neoformans* infection: Pulmonary phagocyte subsets from (A) female and (B) male Balb/c mice were identified by surface staining with the antibodies described in the methods. Cells were stained for surface GLUT-1 and intracellular HK, G6PD, ACAC, and IDH2, followed by flow cytometric analysis. The percentage of cells positive for each metabolic marker was determined within each pulmonary phagocyte subsets. Data are presented as mean (SEM) from 2 independent experiments, with 4 pooled mice per group in each experiment.

Female mice showed trends toward decreased G6PD and ACAC expression following infection, although these differences did not reach statistical significance (p=0.07; Fig. 6A) No corresponding changes were observed in male mice (Fig. 6B). IDH2 expression was significantly increased in AMs from male mice following *C. neoformans* infection (Fig. 6B), whereas no significant changes were observed in female mice (Fig. 6A). The functional significance of this increase remains unclear and requires further investigation. Additionally, no statistically significant changes in ATP5a, ASS1 or PRDX2 expression were detected in the pulmonary phagocyte subsets examined in either sex (Fig. S1 and S2).

Collectively, these findings indicate that pulmonary phagocytes exhibit subset dependent metabolic responses to cryptococcal infection, with the most prominent changes in glycolytic markers occurring in the antifungal female Ly6c^-^ monocyte-like macrophages.

## DISCUSSION

The complexity of immune cells is determined not only by intricate signaling pathways but also by the interconnected metabolic processes that regulate their functions [25]. The overall efficiency of the immune response is highly dependent on the metabolic state of immune cells, which directly influences their activation and antimicrobial activity [26]. Thus, many recent studies have been directed toward understanding the metabolic interactions between immune cells and pathogens.

The present study demonstrates that *C. neoformans* induces coordinated metabolic reprogramming in J774A.1 macrophages. Early increases in GLUT-1, HK and HIF-1α, together with increased lactate production, support enhanced glycolytic metabolism. The induction of GLUT-1 and HK agrees with the established immunometabolic response of activated macrophages, in which increased glucose uptake and phosphorylation support rapid ATP production and antimicrobial functions [27]. Although the cells in this study are not activated to a particular phenotype at this stage, our data indicate that these cells are likely moving towards M1 activation. HIF-1α is a major regulator and promotes protective glycolytic and inflammatory responses during bacterial and fungal infections [28]. The increased HIF-1α observed here may therefore contribute to the induction of GLUT-1 and HK. Increased intracellular lactate is also compatible with greater conversion of pyruvate to lactate. Collectively, these findings suggest a Warburg effect, but further studies are needed to understand if lactate is directly inhibiting *Cryptococcus* or if lactate could directly enhance other antifungal mechanisms like stimulating levels of ROS, NO, or cytokines.

We observed a reduction in ATP5a and mitochondrial IDH2, suggesting a reduced dependence on TCA cycle and OXPHOS. Our mitochondrial results are consistent with previous studies of cryptococcal infection. Studies have shown that *C. neoformans* causes mitochondrial depolarization, mitochondrial fission, and activation of macrophage stress pathways [29]. More recently, a study showed that metabolically active *C. neoformans* increased macrophage glycolysis and altered mitochondrial activity, whereas dormant organisms produced a different metabolic response [30]. Our observations identified changes in proteins involved in ATP synthesis and mitochondrial isocitrate metabolism but additional studies focusing on enzyme activity or respiratory flux are required to further confirm impaired OXPHOS.

Our study also showed that cryptococcal infection of antifungal J774A.1 macrophages led to reduced G6PD and PRDX2 protein expression, indicating broader changes in NADPH production and antioxidant defense. Despite the reduction in G6PD expression, H99 treated macrophages showed increased intracellular ROS and higher *Nox2* and *p47phox* mRNA expression. Reduced G6PD does not establish a loss of PPP activity or NADPH availability; residual NADPH could still support ROS production. Thus, further studies involving measurements of PPP flux, NADPH levels, and NOX2 activity will be needed to determine whether residual PPP-derived NADPH supports the antifungal ROS response to H99. Consistent with a role for the PPP in this response, 6-AN treatment reduced both measured ROS and *Nox2/p47phox* expression. In neutrophils, increased PPP flux supports the oxidative burst and microbial killing [31]. However, in our macrophage model, 6-AN treatment reduced cryptococcal CFUs. One possible explanation is that PPP inhibition limited NADPH dependent glutathione and thioredoxin recycling (GSH/GSSH), thereby increasing oxidative pressure within infected macrophages. Additionally, reduced PRDX2 expression may reinforce this redox environment. Thus, our findings only suggest that PPP inhibition enhances antifungal activity, although the underlying macrophage-specific mechanism focusing on measuring levels of NADPH and GSH/GSSH requires further investigation.

We also observed decreased protein expression of ACAC and ASS1. Reduced ACAC suggests decreased de novo fatty-acid synthesis, which may affect membrane remodeling and inflammatory signaling. Studies have shown that fatty acid use by macrophages during cryptococcal infection is known to vary with fungal metabolic state [30] and *C. neoformans* can incorporate macrophage derived lipids through phospholipase B1 dependent mechanisms [32]. Thus, reduced ACAC may therefore reflect either a host response that restricts lipid availability or pathogen driven remodeling of intracellular niche. Similarly, reduced ASS1 could limit regeneration of arginine from citrulline and therefore can affect NO production. ASS1 dependent arginine recycling has shown to support the control of other intracellular pathogens [33]. Thus, direct measurements of lipid synthesis, arginine metabolites, NOS2 activity and NO will be required to determine the functional consequences of these changes.

Finally, our *in vivo* results have shown increased GLUT-1 and HK expression in female Ly6c^-^monocyte-like macrophages which is consistent with our previously described antifungal activity [11]. However, other metabolic markers did not separate antifungal and permissive populations as clearly. This could be due to several reasons. Cytokines, hormones, nutrient availability, differentiation state and interactions among pulmonary cells may explain why the *in vitro* pattern was only partially reproduced *in vivo*. Additionally, the *in vitro* expression of individual markers was assessed at different timepoints based on their expected activation kinetics, whereas all *in vivo* studies were examined at a single time point: 3 days after infection. Therefore, this time point may not have been optimal for detecting changes across all metabolic pathways, as some responses may occur earlier or require later stages of infection to become apparent.

Combining all this, we have proposed a schematic diagram explaining the antifungal metabolic and redox responses in J774A.1 macrophages following cryptococcal uptake (Fig. 7). In summary, *C. neoformans* induced an early glycolytic response in antifungal J774A.1 macrophages followed by reduced expression of proteins associated with mitochondrial respiration, PPP, lipid synthesis, arginine metabolism, and peroxide detoxification. This study suggests that distribution of glucose-derived carbon may influence the outcome of macrophage- *Cryptococcus* interactions. Establishing the pathways and their cell specific roles may identify metabolic targets that may strengthen antifungal immunity. Current studies in our laboratory are further investigating these pathways by measuring the activity of key metabolic enzymes and evaluating the expression of pathway-associated genes using real-time PCR.

**Fig 7:**
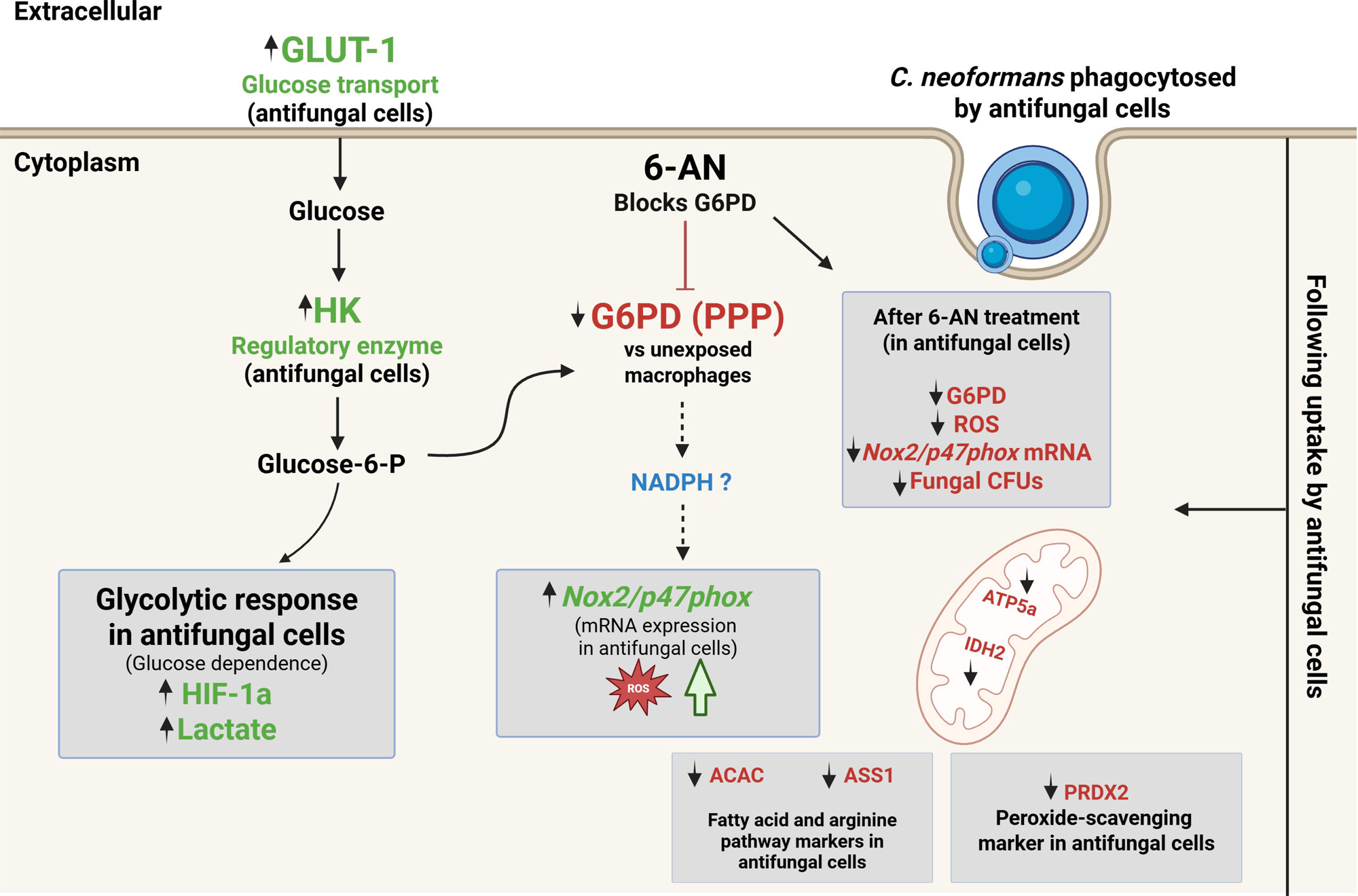
Schematic diagram showing metabolic and redox response of antifungal J774A.1 macrophages following *C. neoformans* uptake. Cryptococcal antifungal activity was associated with increased GLUT-1 and HK expression, glucose dependence, HIF-1α expression and lactate accumulation. G6PD positive cells decreased relative to unexposed macrophages, whereas intracellular ROS and *Nox2/p47phox* mRNA expression increased. Uptake of *C. neoformans* by antifungal cells was also associated with reduced ATP5a, IDH2, ACAC, ASS1, and PRDX2 expression. In *Cryptococcus*-exposed antifungal macrophages, 6-AN treatment further reduced the percentage of G6PD positive cells and decreased ROS, *Nox2/p47phox* mRNA expression, and fungal CFUs relative to vehicle treatment. (Green labels indicate increased expression while red labels indicate decreased expression. Dashed arrows indicate proposed connections that need further investigation such as PPP flux, NADPH activity, and NOX2 activity.)

## SUPPLEMENTARY FIGURES

**Fig S1: Expression of proteins involved in OXPHOS, arginine biosynthesis, and cell redox homeostasis pathway in female pulmonary phagocyte subsets during *C. neoformans* infection:** Pulmonary phagocyte subsets from female Balb/c mice were identified by surface staining with the antibodies described in the methods. Cells were additionally stained intracellularly for ATP5a, PRDX2, and ASS1, followed by flow cytometric analysis. The percentages of cells positive for these metabolic markers were determined by flow cytometry in female pulmonary phagocyte subsets. Data are presented as mean (SEM) from 2 independent experiments, with 4 pooled mice per group in each experiment.

**Fig S2: Expression of proteins involved in OXPHOS, arginine biosynthesis, and cell redox homeostasis pathway in male pulmonary phagocyte subsets during *C. neoformans* infection:** Pulmonary phagocyte subsets from male Balb/c mice were identified by surface staining with the antibodies described in the methods. Cells were additionally stained intracellularly for ATP5a, PRDX2, and ASS1, followed by flow cytometric analysis. The percentages of cells positive for these metabolic markers were determined by flow cytometry in male pulmonary phagocyte subsets. Data are presented as mean (SEM) from 2 independent experiments, with 4 pooled mice per group in each experiment.

**Fig S3: Gating strategy for identification of murine pulmonary phagocyte populations.** Gates were established using single color compensation controls, along with unstained, viability dye controls, and isotype controls. Pulmonary leukocytes were identified as CD45^+^ cells. Following leukocyte gating, AMs were defined as CD11b^-^, and CD11c^hi^; CD11b^+^DCs were defined as CD11c^+^, CD11b^int^; CD103^+^ DCs were defined as CD11c^+^, CD103^+^; interstitial macrophages were defined as CD11c^-^, F4/80^+^; Ly6C^+^ monocyte-like macrophages were defined as CD11c^-^, MHC II^-^, Ly6C^+^; and Ly6C^-^ monocyte-like macrophages were defined as CD11c^-^, MHC II^-^, Ly6C^-^. Each identified population was subsequently analyzed for the expression of the respective metabolic marker.

**Fig S4: Glycolysis inhibition does not alter intracellular ROS levels in *C. neoformans* treated J774A.1 macrophages.** J774A.1 macrophages were incubated with *C. neoformans* KN99 mCherry strain at a ratio of 20:1 for 2h. Cells were then treated with 2-deoxy glucose for 30 mins, followed by CellROX reagent for 30 mins. Following treatment, cells were treated with CellRox reagent for 30 mins. Intracellular ROS was measured by flow cytometry. Data shown are means ± standard errors of the means (SEM) from three independent experiments (n=3).

## REFERENCES

1. Aslanyan, L.; Sanchez, D.A.; Valdebenito, S.; Eugenin, E.A.; Ramos, R.L.; Martinez, L.R. The Crucial Role of Biofilms in Cryptococcus Neoformans Survival within Macrophages and Colonization of the Central Nervous System. J Fungi (Basel) 2017, 3, 10, doi:10.3390/jof3010010.

2. Rajasingham, R.; Govender, N.P.; Jordan, A.; Loyse, A.; Shroufi, A.; Denning, D.W.; Meya, D.B.; Chiller, T.M.; Boulware, D.R. The Global Burden of HIV-Associated Cryptococcal Infection in Adults in 2020: A Modelling Analysis. Lancet Infect Dis 2022, 22, 1748–1755, doi:10.1016/S1473-3099(22)00499-6.

3. Heung, L.J.; Hohl, T.M. Inflammatory Monocytes Are Detrimental to the Host Immune Response during Acute Infection with Cryptococcus Neoformans. PLoS Pathog 2019, 15, e1007627, doi:10.1371/journal.ppat.1007627.

4. Paiva, A.L.C.; Aguiar, G.B. de; Lovato, R.M.; Zanetti, A.V.D.; Panagopoulos, A.T.; Veiga, J.C.E. Cryptococcoma Mimicking a Brain Tumor in an Immunocompetent Patient: Case Report of an Extremely Rare Presentation. Sao Paulo Med J 2018, 136, 492–496, doi:10.1590/1516-3180.2017.0046210417.

5. Velamakanni, S.S.; Bahr, N.C.; Musubire, A.K.; Boulware, D.R.; Rhein, J.; Nabeta, H.W. Central Nervous System Cryptococcoma in a Ugandan Patient with Human Immunodeficiency Virus. Med Mycol Case Rep 2014, 6, 10–13, doi:10.1016/j.mmcr.2014.08.003.

6. Hawkins, A.N.; Determann, B.F.; Nelson, B.N.; Wozniak, K.L. Transcriptional Changes in Pulmonary Phagocyte Subsets Dictate the Outcome Following Interaction With The Fungal Pathogen Cryptococcus Neoformans. Front Immunol 2021, 12, 722500, doi:10.3389/fimmu.2021.722500.

7. Pellon, A.; Begum, N.; Sadeghi Nasab, S.D.; Harzandi, A.; Shoaie, S.; Moyes, D.L. Role of Cellular Metabolism during Candida-Host Interactions. Pathogens 2022, 11, 184, doi:10.3390/pathogens11020184.

8. Howard, N.C.; Khader, S.A. Immunometabolism during Mycobacterium Tuberculosis Infection. Trends Microbiol 2020, 28, 832–850, doi:10.1016/j.tim.2020.04.010.

9. Traven, A.; Naderer, T. Central Metabolic Interactions of Immune Cells and Microbes: Prospects for Defeating Infections. EMBO Rep 2019, 20, e47995, doi:10.15252/embr.201947995.

10. Lachmandas, E.; Boutens, L.; Ratter, J.M.; Hijmans, A.; Hooiveld, G.J.; Joosten, L.A.B.; Rodenburg, R.J.; Fransen, J.A.M.; Houtkooper, R.H.; van Crevel, R.;, et al. Microbial Stimulation of Different Toll-like Receptor Signalling Pathways Induces Diverse Metabolic Programmes in Human Monocytes. Nat Microbiol 2016, 2, 16246, doi:10.1038/nmicrobiol.2016.246.

11. Nair, A.S.; Wozniak, K.L. MHC-I Upregulation and Increased Glucose Dependence Define Innate Antifungal Immune Responses to Cryptococcus Neoformans. J Infect Dis 2026, jiag144, doi:10.1093/infdis/jiag144.

12. Gleeson, L.E.; Sheedy, F.J.; Palsson-McDermott, E.M.; Triglia, D.; O’Leary, S.M.; O’Sullivan, M.P.; O’Neill, L.A.J.; Keane, J. Cutting Edge: Mycobacterium Tuberculosis Induces Aerobic Glycolysis in Human Alveolar Macrophages That Is Required for Control of Intracellular Bacillary Replication. J Immunol 2016, 196, 2444–2449, doi:10.4049/jimmunol.1501612.

13. Tucey, T.M.; Verma, J.; Harrison, P.F.; Snelgrove, S.L.; Lo, T.L.; Scherer, A.K.; Barugahare, A.A.; Powell, D.R.; Wheeler, R.T.; Hickey, M.J.;, et al. Glucose Homeostasis Is Important for Immune Cell Viability during Candida Challenge and Host Survival of Systemic Fungal Infection. Cell Metab 2018, 27, 988–1006.e7, doi:10.1016/j.cmet.2018.03.019.

14. Gonçalves, S.M.; Duarte-Oliveira, C.; Campos, C.F.; Aimanianda, V.; Ter Horst, R.; Leite, L.; Mercier, T.; Pereira, P.; Fernández-García, M.; Antunes, D.;, et al. Phagosomal Removal of Fungal Melanin Reprograms Macrophage Metabolism to Promote Antifungal Immunity. Nat Commun 2020, 11, 2282, doi:10.1038/s41467-020-16120-z.

15. Schwartz, L.; Supuran, C.T.; Alfarouk, K.O. The Warburg Effect and the Hallmarks of Cancer. Anticancer Agents Med Chem 2017, 17, 164–170, doi:10.2174/1871520616666161031143301.

16. Rosa, R.L.; Berger, M.; Santi, L.; Driemeier, D.; Barros Terraciano, P.; Campos, A.R.; Guimarães, J.A.; Vainstein, M.H.; Yates, J.R.; Beys-da-Silva, W.O. Proteomics of Rat Lungs Infected by Cryptococcus Gattii Reveals a Potential Warburg-like Effect. J Proteome Res 2019, 18, 3885–3895, doi:10.1021/acs.jproteome.9b00326.

17. Shi, L.; Salamon, H.; Eugenin, E.A.; Pine, R.; Cooper, A.; Gennaro, M.L. Infection with Mycobacterium Tuberculosis Induces the Warburg Effect in Mouse Lungs. Sci Rep 2015, 5, 18176, doi:10.1038/srep18176.

18. Jha, A.K.; Huang, S.C.-C.; Sergushichev, A.; Lampropoulou, V.; Ivanova, Y.; Loginicheva, E.; Chmielewski, K.; Stewart, K.M.; Ashall, J.; Everts, B.;, et al. Network Integration of Parallel Metabolic and Transcriptional Data Reveals Metabolic Modules That Regulate Macrophage Polarization. Immunity 2015, 42, 419–430, doi:10.1016/j.immuni.2015.02.005.

19. Tannahill, G.M.; Curtis, A.M.; Adamik, J.; Palsson-McDermott, E.M.; McGettrick, A.F.; Goel, G.; Frezza, C.; Bernard, N.J.; Kelly, B.; Foley, N.H.;, et al. Succinate Is an Inflammatory Signal That Induces IL-1β through HIF-1α. Nature 2013, 496, 238–242, doi:10.1038/nature11986.

20. Batista-Gonzalez, A.; Vidal, R.; Criollo, A.; Carreño, L.J. New Insights on the Role of Lipid Metabolism in the Metabolic Reprogramming of Macrophages. Front Immunol 2019, 10, 2993, doi:10.3389/fimmu.2019.02993.

21. Ahl, P.J.; Hopkins, R.A.; Xiang, W.W.; Au, B.; Kaliaperumal, N.; Fairhurst, A.-M.; Connolly, J.E. Met-Flow, a Strategy for Single-Cell Metabolic Analysis Highlights Dynamic Changes in Immune Subpopulations. Commun Biol 2020, 3, 305, doi:10.1038/s42003-020-1027-9.

22. Erlich, J.R.; To, E.E.; Luong, R.; Liong, F.; Liong, S.; Oseghale, O.; Miles, M.A.; Bozinovski, S.; Brooks, R.D.; Vlahos, R.;, et al. Glycolysis and the Pentose Phosphate Pathway Promote LPS-Induced NOX2 Oxidase- and IFN-β-Dependent Inflammation in Macrophages. Antioxidants (Basel) 2022, 11, 1488, doi:10.3390/antiox11081488.

23. Pouysségur, J.; Marchiq, I.; Parks, S.K.; Durivault, J.; Ždralević, M.; Vucetic, M. “Warburg Effect” Controls Tumor Growth, Bacterial, Viral Infections and Immunity - Genetic Deconstruction and Therapeutic Perspectives. Semin Cancer Biol 2022, 86, 334–346, doi:10.1016/j.semcancer.2022.07.004.

24. Bamgbose, T.T.; Igiehon, O.O.; Nion-Fieira, J.; Palmer, C.R.; Andriichuk, A.; Woolard, M.D. Fatty Acid Metabolism and Lipid Channeling in Macrophages: Mechanisms of Inflammation, Resolution, and Lipotoxicity. Front Immunol 2026, 17, 1839069, doi:10.3389/fimmu.2026.1839069.

25. Xu, R.; He, X.; Xu, J.; Yu, G.; Wu, Y. Immunometabolism: Signaling Pathways, Homeostasis, and Therapeutic Targets. MedComm (2020) 2024, 5, e789, doi:10.1002/mco2.789.

26. Makowski, L.; Chaib, M.; Rathmell, J.C. Immunometabolism: From Basic Mechanisms to Translation. Immunol Rev 2020, 295, 5–14, doi:10.1111/imr.12858.

27. Shao, C.; Lin, S.; Liu, S.; Jin, P.; Lu, W.; Li, N.; Zhang, Y.; Bo, L.; Bian, J. HIF1α-Induced Glycolysis in Macrophage Is Essential for the Protective Effect of Ouabain during Endotoxemia. Oxid Med Cell Longev 2019, 2019, 7136585, doi:10.1155/2019/7136585.

28. Li, C.; Wang, Y.; Li, Y.; Yu, Q.; Jin, X.; Wang, X.; Jia, A.; Hu, Y.; Han, L.; Wang, J.;, et al. HIF1α-Dependent Glycolysis Promotes Macrophage Functional Activities in Protecting against Bacterial and Fungal Infection. Sci Rep 2018, 8, 3603, doi:10.1038/s41598-018-22039-9.

29. Coelho, C.; Souza, A.C.O.; Derengowski, L. da S.; de Leon-Rodriguez, C.; Wang, B.; Leon-Rivera, R.; Bocca, A.L.; Gonçalves, T.; Casadevall, A. Macrophage Mitochondrial and Stress Response to Ingestion of Cryptococcus Neoformans. J Immunol 2015, 194, 2345–2357, doi:10.4049/jimmunol.1402350.

30. Marina, C.L.; de Castro, R.J.A.; Bellozi, P.; Cruz, A.M.; Bürgel, P.H.; Potter, P.G.W.; Beall, C.; Tavares, A.H.; De Bem, A.; Alanio, A.;, et al. Immunometabolic Reprogramming in Macrophages Infected with Active and Dormant Cryptococcus Neoformans: Differential Modulation of Respiration, Glycolysis, and Fatty Acid Utilization. Infect Immun 2025, 93, e0048724, doi:10.1128/iai.00487-24.

31. Britt, E.C.; Lika, J.; Giese, M.A.; Schoen, T.J.; Seim, G.L.; Huang, Z.; Lee, P.Y.; Huttenlocher, A.; Fan, J. Switching to the Cyclic Pentose Phosphate Pathway Powers the Oxidative Burst in Activated Neutrophils. Nat Metab 2022, 4, 389–403, doi:10.1038/s42255-022-00550-8.

32. Wright, L.C.; Santangelo, R.M.; Ganendren, R.; Payne, J.; Djordjevic, J.T.; Sorrell, T.C. Cryptococcal Lipid Metabolism: Phospholipase B1 Is Implicated in Transcellular Metabolism of Macrophage-Derived Lipids. Eukaryot Cell 2007, 6, 37–47, doi:10.1128/EC.00262-06.

33. Qualls, J.E.; Subramanian, C.; Rafi, W.; Smith, A.M.; Balouzian, L.; DeFreitas, A.A.; Shirey, K.A.; Reutterer, B.; Kernbauer, E.; Stockinger, S.;, et al. Sustained Generation of Nitric Oxide and Control of Mycobacterial Infection Requires Argininosuccinate Synthase 1. Cell Host Microbe 2012, 12, 313–323, doi:10.1016/j.chom.2012.07.012.

