## Supplementary figures and images for "Host Metabolic Reprogramming Defines Macrophage Antifungal Activity Against *Cryptococcus neoformans*"

### Supplementary Figure 1

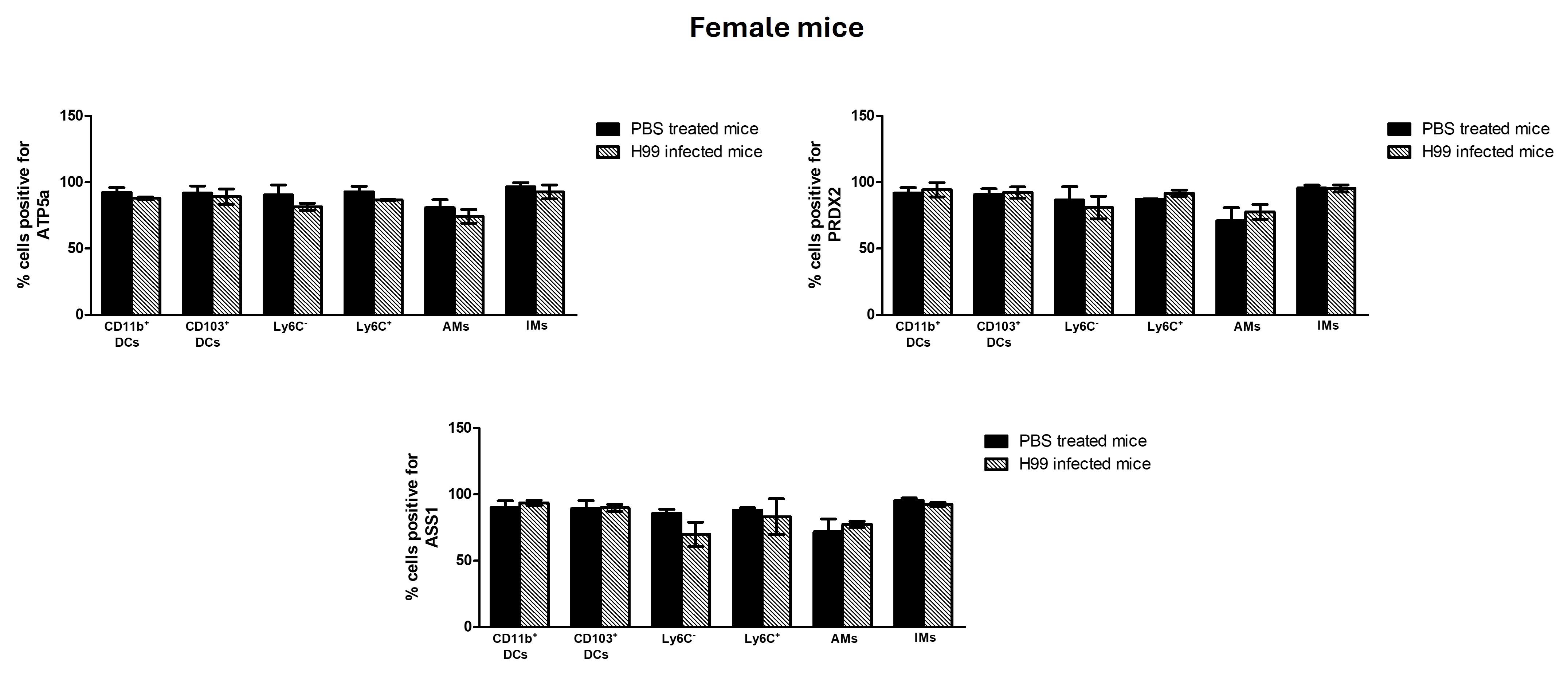

### Supplementary Figure 2

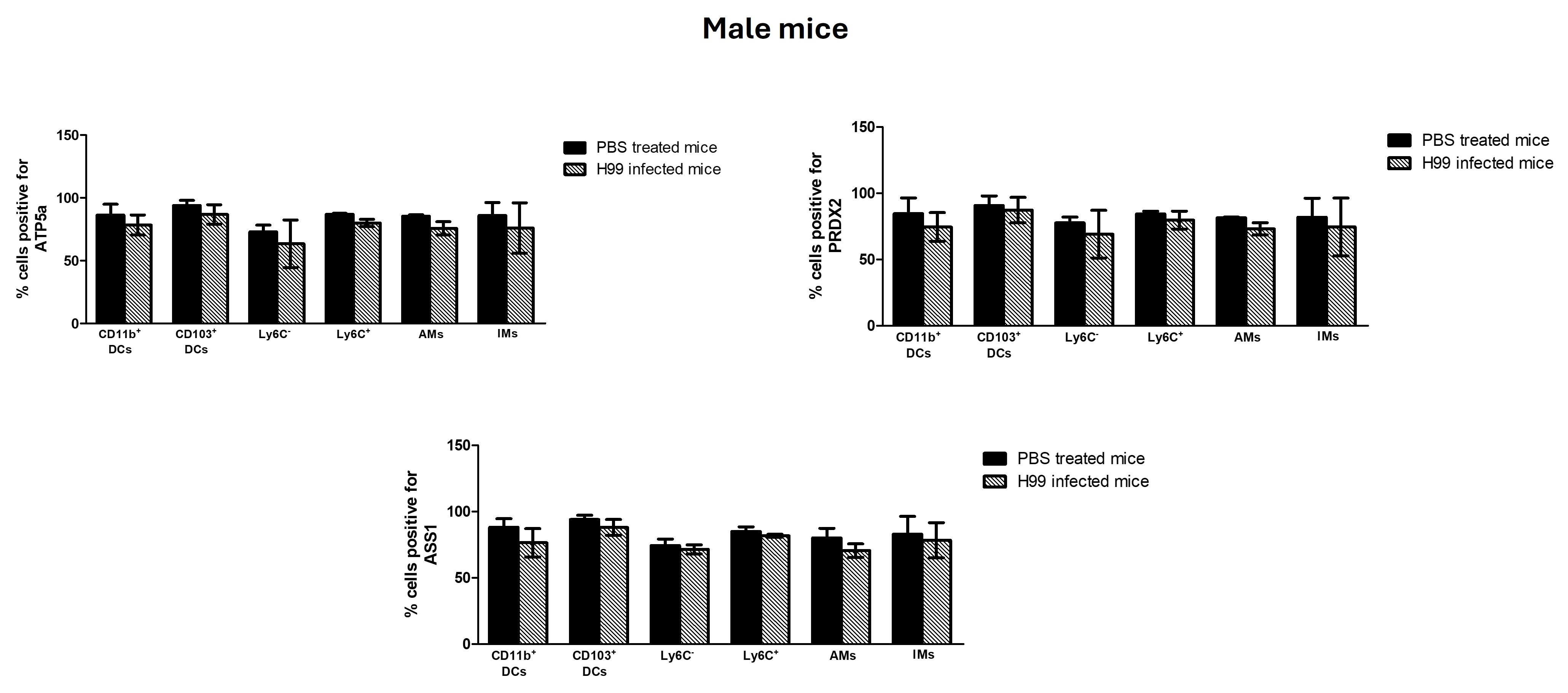

### Supplementary Figure 3

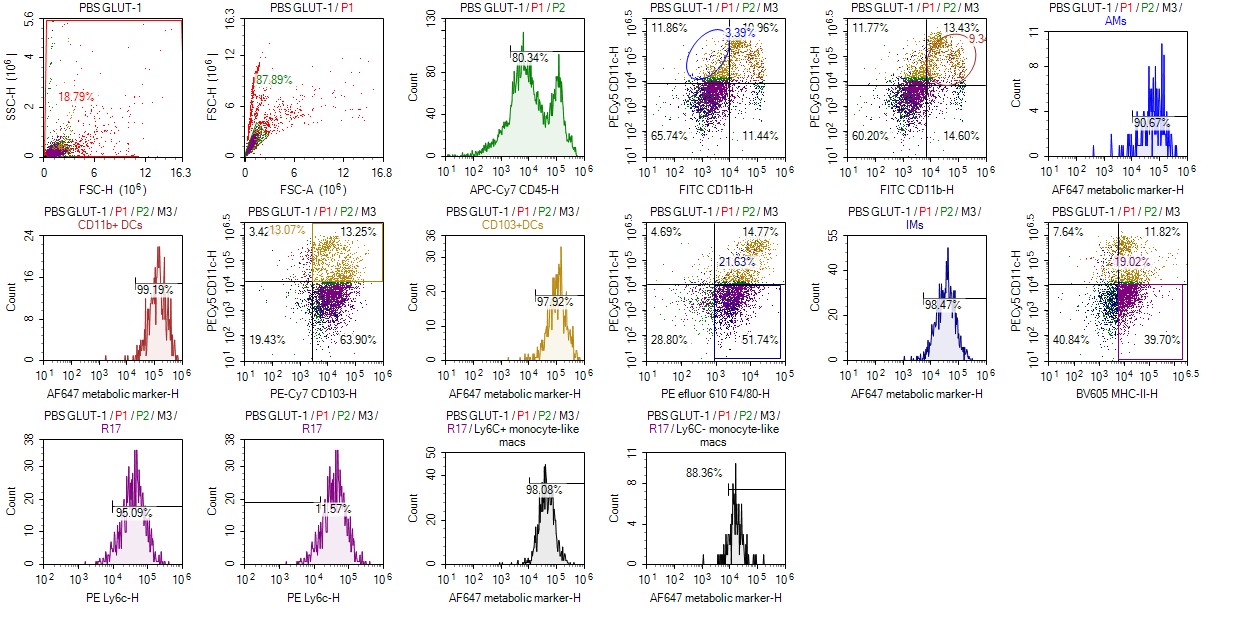

### Supplementary Figure 4

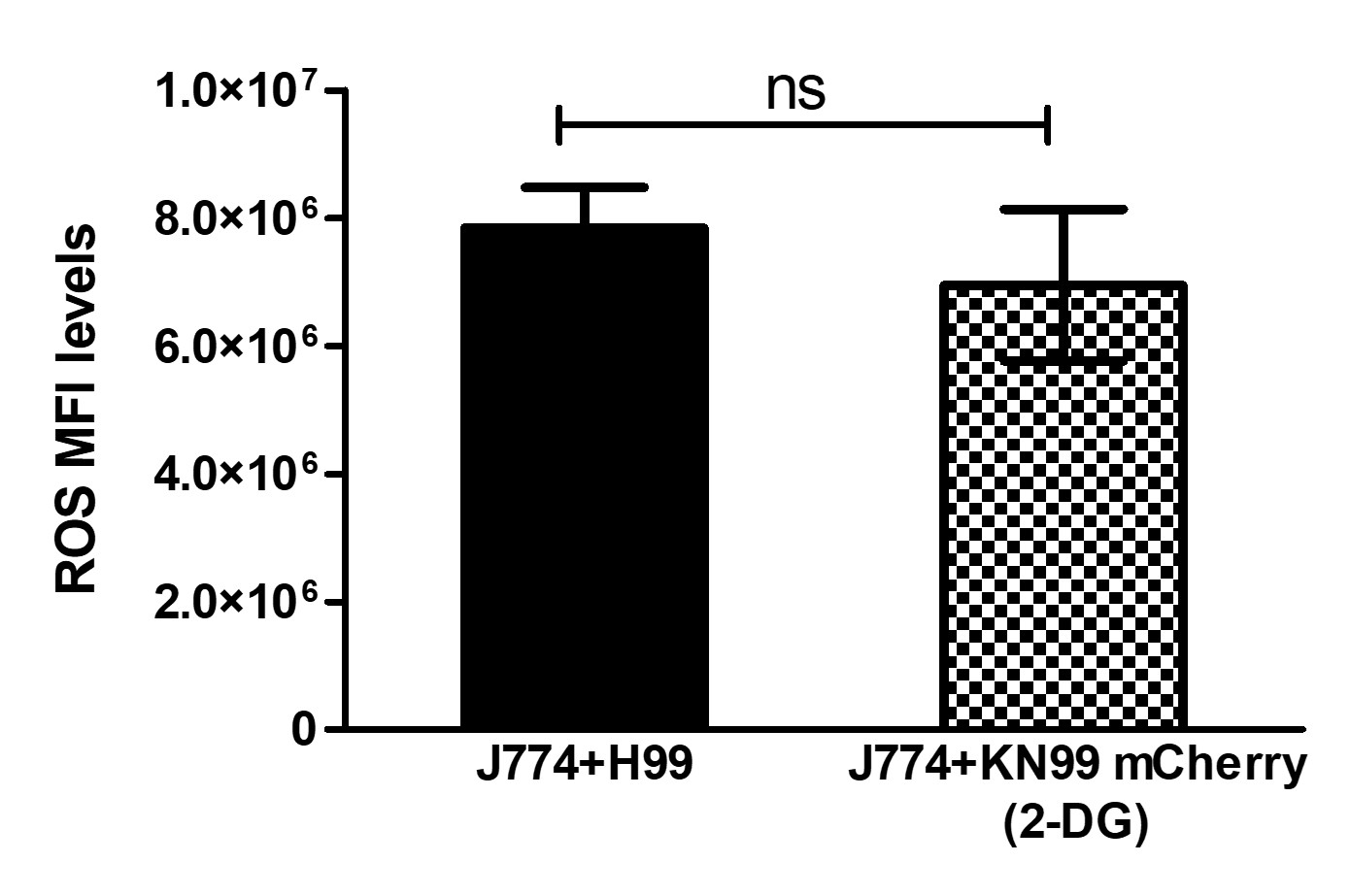
